# Structural insights into the initiation of free radical formation in the Class Ib ribonucleotide reductases

**DOI:** 10.1101/2023.06.06.543845

**Authors:** Lumbini R. Yadav, Shekhar C. Mande

**Affiliations:** National Centre for Cell Science, SPPU Campus, Ganeshkhind, Pune- 411007, INDIA and; Bioinformatics Centre, Savitribai Phule Pune University, Ganeshkhind, Pune- 411007

**Keywords:** RNR’s-Ribonucleotide reductase, Mtb-*Mycobacterium tuberculosis*, Mth-*Mycobacterium thermoresistibile*, Tyr- free radical, NrdF2I- NrdF2:NrdI complex

## Abstract

Class I ribonucleotide reductases consisting of R1 and R2 subunits convert ribonucleoside diphosphates to deoxyribonucleoside diphosphates involving an intricate free radical mechanism. The generation of free radicals in the Class Ib ribonucleotide reductases is mediated by reaction involving di-manganese ions in the R2 subunits and is externally assisted by flavodoxin-like NrdI subunit. This is unlike Class Ia ribonucleotide reductases, where the free radical generation is initiated at its di-iron centre in the R2 subunits with no external support from another subunit. Despite much work on the R2 subunits of Class Ib ribonucleotide reductases, also referred as NrdF, and its partner NrdI, the structural details of free radical generation remain largely unknown. In this study we have determined the crystal structures of Mycobacterial NrdI in oxidized and reduced forms, and similarly those of NrdF2: NrdI complex (NrdF2I). These structures provide the first atomic view of the mechanism of free radical generation in the R2 subunit. We propose that oxygen molecule accesses FMN through a well-formed channel in NrdI, seen clearly in the crystal structure, and upon electron transfer is converted to a superoxide ion. Similarly, a path for superoxide radical transfer between NrdI and NrdF2 is also observed. A delocalised Mn ion in the R2 subunit is seen in the electron density, which attacks Tyr 110 to produce a Tyr• free radical. Finally, a solvent channel to the dimanganese-binding site is observed to complete the cycle. The study therefore provides important structural clues on the initiation of free radical generation in the R2 subunit of the ribonucleotide reductase complex.

**Significance statement:** Ribonucleotide reductases generate the deoxyribonucleotide pool in the cell for DNA replication and repair. The enzymes utilise free radical mechanism, where the mechanism of radical formation defines different classes of ribonucleotide reductases. Class Ib ribonucleotide reductases generate the free radical though di-manganese chemistry, assisted externally by NrdI. We describe here structural features required to achieve this mechanism. The structures clearly show a tunnel for oxygen access to the FMN site, tunnel to transport the consequent superoxide radical that is formed, a dislocated activated Mn, which appears to coordinate with a Tyrosine residue to form Tyr• radical and a water channel to complete the reaction cycle, thus enhancing our understanding of the steps of free radical generation.

## 1. Introduction

Reduction of ribonucleoside di-/tri phosphates into deoxyribonucleoside di-/tri phosphates is one of the most fundamental biochemical steps in all life forms^1^. Ribonucleotide reductases (RNR’s), which catalyse this step, use a unique free radical mechanism for the conversion. RNR’s have been classified into three distinct classes depending upon the cofactor they use, and the radicals that mediate the catalytic reaction. Among the RNR’s, Class I RNR’s typically consist of two subunits, R1 and R2, where the R1 subunits mediate the catalytic conversion of ribonucleosides into deoxyribonucloesides, whereas free radical generation and maintenance happens in the R2 subunits^2^. The Class I ribonucleotide reductases have further been classified into Class Ia-Ie, based on their allosteric regulation and accessory proteins they utilize^3,4^. Class Ia RNR R2 subunits use Fe to facilitate generation of Tyr• radical in presence of oxygen, while Class Ib R2 subunits need external assistance from NrdI protein, and generate the Tyr• radical mediated by Manganese, and Oxygen^2^. Large numbers of elegant biochemical and structural studies have revealed different aspects of free radical transfer and utilization during the enzymatic cycle^5,6,7^. Moreover, structural studies have also been important in enhancing our understanding of the allosteric regulation mechanisms in different RNR’s ^8,9,10^.

Mycobacteria possess two RNR classes, but only Class Ib RNR has been shown to be essential for growth in *M. tuberculosis*^11^. Class Ib RNR in Mycobacteria have therefore been considered to be attractive drug targets. Class Ib genes in prokaryotes are encoded by the *nrdE*, *nrdF* and *nrdI* subunits, with other accessory genes such as the *nrdH*. The *nrdE* and *nrdF* genes in Class Ib RNR’s and their homologs *nrdA* and *nrdB* in Class Ia RNR’s are more generally referred to as the R1 and R2 subunits. Although the catalytic cycle of conversion of ribonucleoside di-phosphate to deoxyribonucleoside di-phosphate happens in the NrdE subunits of Class Ib RNR’s, free radical generation using atmospheric oxygen occurs at the di-metal binding site of the NrdF assisted by the NrdI subunit.

Much work using structural, biochemical and spectroscopic studies has been carried out to enhance our understanding of the Class Ia RNR’s, especially that related to radical transfer over a long range between R1 and R2, and mode of regulation by nucleotides. Mechanisms of these are likely to be similar in Class Ib RNR’s^12,9,13,14^. O_2_^•-^ mediated assembly of Mn^III^ Mn^IV^ intermediate in Class Ib RNR and consequent Tyr• radical formation has also been observed experimentally^15^. Nonetheless, understanding of free radical generation by the di-Manganese centre assisted by NrdI in the Class Ib RNR’s is incomplete. We present here crystal structures of NrdI in oxidized and reduced forms at a resolution of 1.2Å, and those of NrdF2I complex at a resolution of 3Å. The structural analysis clearly outlines the early steps in free radical generation in the Class Ib RNR family.

## 2. Results

### 2.1. Overall structural analysis of NrdI

The structure of *Mycobacterium thermoresistibile* (Mth) NrdI was determined at 1.2Å resolution in its oxidised and reduced forms (Supplementary Table1). In both the structures, the last three residues of the sequence and the first two residues are not visible in electron density. Moreover, electron density in the loops near the FMN binding site was not uniformly present. The structure of NrdI is similar to other reported NrdI structures with canonical flavodoxin-like fold (Figure1A). The root mean square deviation (RMSD) among the available structures of NrdI reveals comparable conformational features among all the structures (Supplementary Table 2).

Electron density for FMN is present at its canonical binding site in the pockets formed by 40s, 50s and 80s loops termed according to flavodoxin (FLDs) nomenclature^16^ (Figure 1A). The FMN isoalloxazine ring is bound in a pocket through hydrogen bonds between main chain atoms of the protein, namely N3 with carbonyl oxygen of Glu 95, O2 with main chain peptidyl nitrogens of Asn 89 and Ala 97, and N5 with peptidyl nitrogen of Gly 51 (in the oxidized form) and carbonyl oxygen of Gly 50 (in the reduced form). The ribitol group also forms three hydrogen bonds involving water and backbone chain atoms of Thr 48. The phosphate group of FMN is anchored by helix dipole of helix 1 and by the main chain nitrogen of the loop present before the helix.

**Figure 1.**
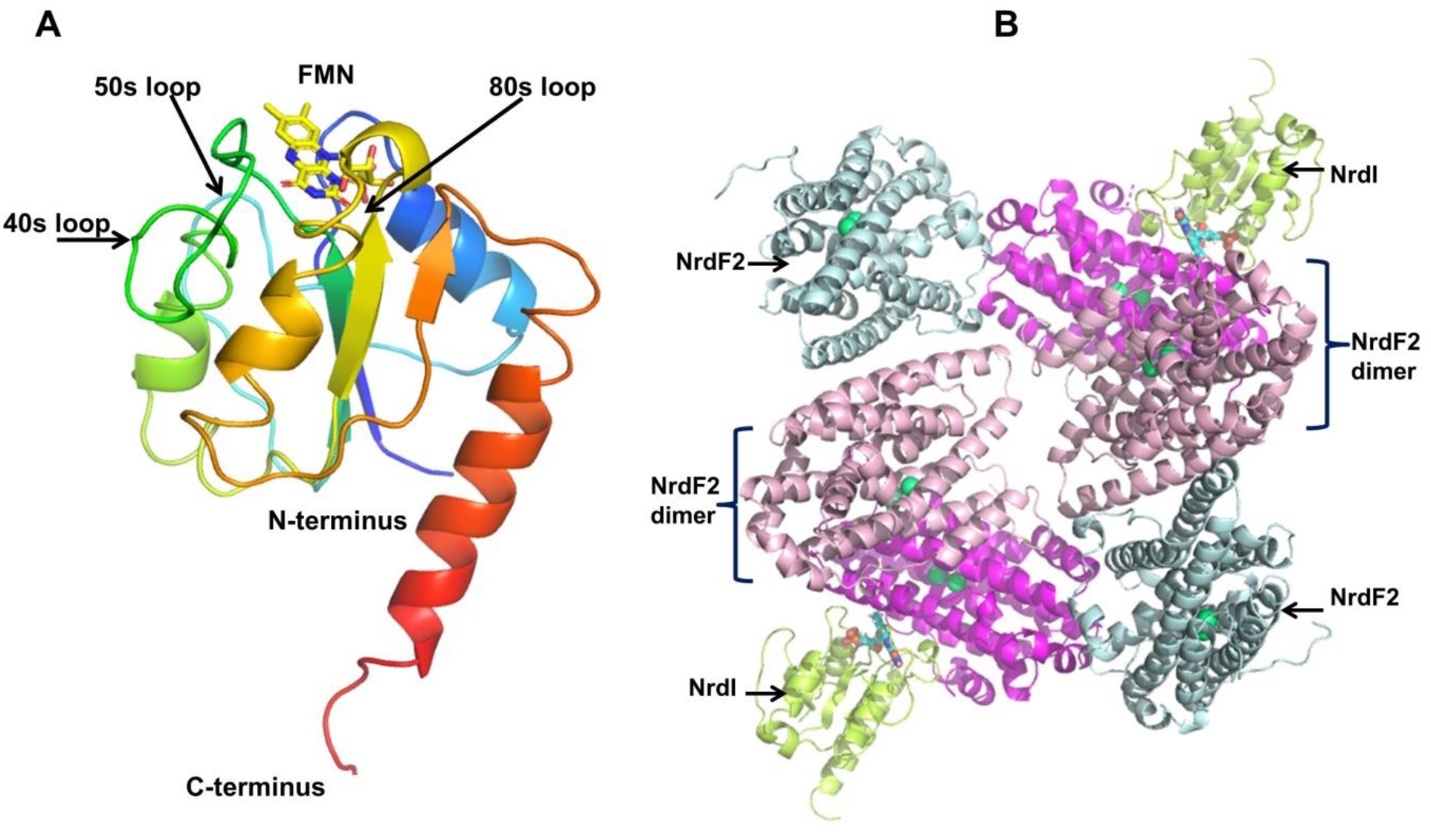
Overview of structure of Mth NrdI and Mtb NrdF2I complex: **(A)** The overall structure of Mth NrdI shown in cartoon representation. The chain has been colour-coded from blue to red starting from N-terminal to C-terminal. The important features of the structure, namely the 50s loop, 40s loop and 80s loop are indicated. These loops not only make important contacts with the FMN moiety, but also participate in association with NrdF2. **(B)** Structure of MtbNrdF2I complex: 6 chains of NrdF2 and 2 of NrdI were observed in the crystal asymmetric unit. The stoichiometry of the complex was such that two chains of the dimer of NrdF interacted with one chain of NrdI. NrdF chain which is not part of physiological dimer colored as palecyan. The functional part of asymmetric unit which is dimer of NrdF2 is colored as hotpink and lightpink while a single NrdI monomer is colored as limongreen. Bound metal ions in NrdF2 are shown in limegreen.

The isoalloxazine ring of FMN is stacked between residues Phe 92 on one side and Tyr 49 on the other. Phe 92 demonstrates a parallel displaced π-π interaction while Tyr 49 shows parallel staggered interaction with FMN^17,18^. The stacked residues phenylalanine and tyrosine are present at a distance of 3.6Å and 4.3Å respectively from the FMN moiety. Tyr 49 is conserved in all the reported NrdI structures, while phenylalanine or tryptophan residues are observed at position-92. These nonbonded π-π interactions are also observed in other NrdI structures and play a role in stabilisation of the isoalloxazine ring of FMN in binding pocket, electron transfer during reduction, and may also have a role in promoting interaction responsible for molecular assembly with NrdF2.

#### 2.1.2. Comparative analysis of Mth NrdI (reduced/oxidised) and other reported structures

The structures of oxidized and reduced forms of NrdI superpose with an overall RMSD of 0.5Å. The closest structural homolog of Mth and *Mycobacterium. tuberculosis* (Mtb) NrdI is *E.coli* NrdI. Major structural differences between the *E. coli* and Mth structures is confined to the N- and C- termini and in the region near the 50s loop. The 50s loop of Mtb NrdI is longer by about 7 residues than that of *E. coli* NrdI (Supplementary Figure 1A). The length of 50s loop in *Aerococous urinae* (PDB ID 6EBQ) is similar to that in Mth and Mtb (Supplementary Figure 1B).

The major conformational change observed in the structures is around the isoalloxazine ring, where a peptide flip at Gly 50 is seen in the reduced form, similar to that observed in other reported structures of NrdI^19^. The peptide flip leads to the formation of a hydrogen bond between carbonyl oxygen of Gly 50 and N5 atom of FMN (Supplementary Figure 1C). Possibly as a consequence of this stabilization, the density of the 50s loop was clearly visible in reduced form unlike that in the oxidised form (Supplementary Figure 2A). B-factors of the main chain in this region also clearly show that 50s loop gets better ordered in the reduced than in the oxidized NrdI (Supplementary Figure 2B).

### 2.2. Structural analysis of Mtb NrdF2I complex

The structures of Mtb NrdF2I complexes in reduced and oxidised forms were determined at ∼3Å resolution (Supplementary Table1). Interestingly, the crystal asymmetric unit consisted of 6 molecules of NrdF2 and 2 molecules of NrdI, with a 2:1 stoichiometry of NrdF2 and NrdI (Figure 1B). The final refined model of NrdF2I complex consists of residues ∼8-293 in NrdF2 molecule with the 32 C-terminal residues missing because of disorder present in this region. This is similar to reported structures of NrdF2 from other organisms^20,21,22^. NrdI in the complex shows density for amino acid residues ∼7 to 143 with backbone chain density clearly visible in the 50s loop. It is clearly seen from the B-factor plot in both oxidised and reduced complex structures that there are two distinct peaks of disorder in the structure, centred on 40s loop and centred on 50s loop. It is interesting to note that the 40s loop is highly ordered in standalone NrdI structure as opposed to the NrdF2I complex. Upon examination of the standalone NrdI structure it appears the ordering of this loop might be due to crystal packing interaction (Supplementary Figure 2B).

The Mtb NrdF2I complex structures superpose with the reported Class Ib structures with a RMSD of 1.0 and 0.6Å for *B. cereus* (PDB ID 4BMO) and *E. coli* (PDB ID 3N39) respectively and 0.8Å for Class Ie *A. urinae* (PDB ID 7MMP) structure (Supplementary Table 3).

#### 2.2.1. Stoichiometry of interaction

As the crystal asymmetric unit contains six chains of NrdF2 calculation of buried surface area between different NrdF2 monomers indicated the correct physiological dimer among all these chains. The buried surface area between two monomers in the physiological dimer of NrdF2 is ∼2000 Å^2^ and between the chains that do not form physiological dimer is ∼850 Å^2^. The two chains of NrdI associate with two physiological dimers of NrdF2 in the crystal asymmetric unit (Figure1B).

The stoichiometry of interaction in the Mtb NrdF2I complex structure is 2:1, unlike other reported structures where the stoichiometry observed to be 1:1. Examination of crystal packing indicated possibility of a clash if the stoichiometry were 1:1. The position where another NrdI subunit would be placed in such a 1:1 complex is occupied by another NrdF2 subunit in our structure. To probe the true stoichiometry of the NrdF2I complex, we performed additional experiments to address the issue. SEC-MALS analysis suggested stoichiometry of 2:1 (Supplementary Figure 3). Similarly, cross linking experiments also indicated 2:1 stoichiometry (Data not shown). Moreover, low resolution cryo-EM map obtained of Mth. NrdF2I structure also suggests the stoichiometry of the complex to be 2:1 (unpublished results). Therefore, from the SEC-MALS, cryo-EM and crosslinking studies we believe that the stoichiometry of Mtb NrdF2I complex is 2:1 and might not be crystallization artefact.

#### 2.2.2. Interaction interface of complex

In the crystal structure of the NrdF2I complex, the association of the two molecules is such that the FMN cofactor of NrdI and di-Manganese atoms of NrdF2 are in close vicinity. Such a placement makes them apparently poised for the radical transfer reaction (Figure 2A). The buried surface area between NrdF2 and NrdI in both the oxidised and reduced forms is almost similar, that is ∼1000 Å^2^ ^23–25^. The interaction between NrdF2 and NrdI is mostly through hydrogen bonds and hydrophobic interactions.

**Figure 2:**
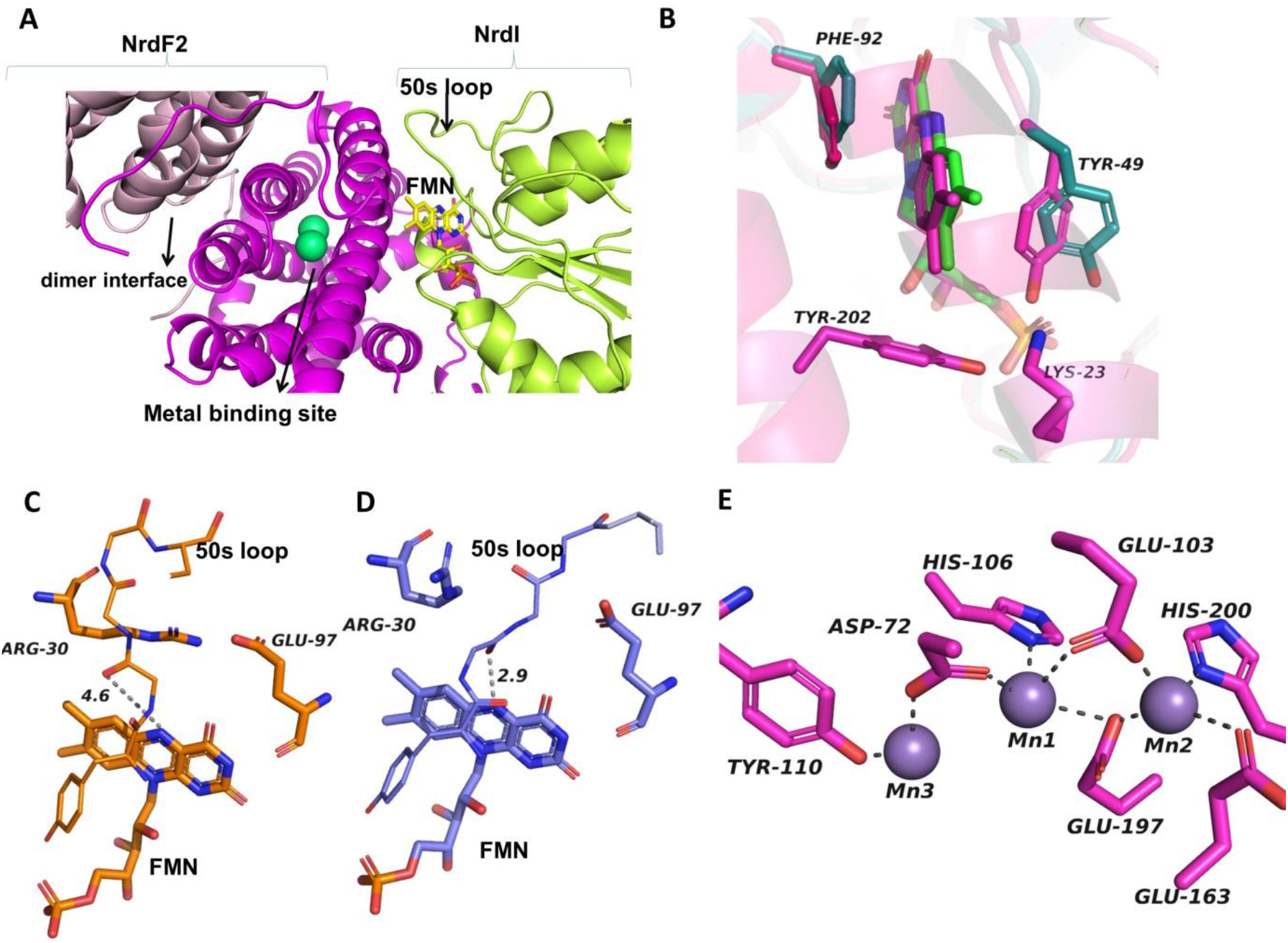
Interface of complex of Mtb NrdF2I: **(A)** Interaction of NrdI and NrdF is characterized by close interactions of the radical transfer groups. FMN of NrdI is placed close to the di-manganese binding site so that superoxide radical produced by FMN can be transported to metal centre. 50s loop of NrdI (green) makes association of NrdF2 (pink) and NrdI more intimate. **(B).** Subtle differences in the structures of NrdI and NrdF2I complex. A rotation in NrdI Tyr 49 side chain of complex was observed promoting closer interactions of NrdF2 and NrdI subunits. **(C)** During complex formation among the structural changes is the reorientation of Arg 30 side chain and 50s loop between the oxidized **(D)** and reduced forms **(E)** Residues involved in Mn binding in the NrdF2I complex.

Superposition of Mth NrdI with that of Mtb NrdF2I complex indicated 90° rotation of Tyr 51 of Mtb NrdI (Mth Tyr 49) towards FMN in the complex. The rotation and movement of the Tyr facilitates its intersubunit interaction with Tyr 202 and Lys 23 of NrdF2 and PO3 group of FMN. The interactions of Lys 23 and Tyr 202 with Tyr 51 appear to stabilise the NrdF2 and NrdI interactions in the complex (Figure 2B).

#### 2.2.3. Comparative analysis of oxidised and reduced Mtb NrdF2I complex

The RMSD of the reported Mtb NrdF2 structure^20^ with our complex structure is 0.35Å indicating no major structural change in NrdF2 in oxidised and reduced forms. The major change between the reduced form of the complex compared to the oxidised form is at interface near the NrdI 50s loop. An ionic interaction is observed between NrdF2-Arg 30 and NrdI-Glu 97 in the oxidised but not in the reduced complex. The orientation of NrdF2-Arg 30 in oxidised complex occludes interaction of 50s loop with FMN but enables its interaction with FMN in the reduced complex. Consequently, in reduced complex the 50s loop reorients and moves towards NrdI facilitating the hydrogen bonding interaction of carbonyl oxygen of Gly 52 and N5 atom of FMN (Figure 2C and 2D).

#### 2.2.4. Interaction at metal site

X-ray fluorescence scan of protein complex crystals indicated the characteristic feature of manganese present in the structure. The peak at ∼6.5 keV corresponds to K edge emission of Mn, thus confirming presence of Mn and not Fe. Residues involved in interaction with metal ion in NrdF2 are His 200, His 106, Glu 163, Glu 103, Glu 197 and Asp72 (Figure 2E). The 2 manganese ions are at a Mn-Mn distance of ∼3.86 Å. Tyr 110 implicated for receiving the free radical is placed close to the di-manganese binding site approximately at a distance of 6Å. Tyr 110 is immersed in a complete hydrophobic environment surrounded by side chains of Phe 65, Ile 190, Ile 193, Phe 171, Leu 68 and Phe 167.

An interesting observation pertains to the orientation of Glu-163 which is one of the Mn-coordinating residues, and whose orientation is different from the equivalent residue Glu-158 from *E.coli*. Among the different NrdF and NrdFI complex structures (with Iron or Manganese) assessed, residues equivalent to Glu-163 are in similar orientation in most structures as that in Mtb NrdF2I complex (Figure 3A). This orientation permits monodentate coordination with Mn ion. On the other hand, in *B.cereus* (4BMU) and *E.coli (3N39)* structures of NrdF2I, the Glu residue exhibits bidentate interaction with manganese^6,26^ (Figure 3B and 3C).

**Figure 3:**
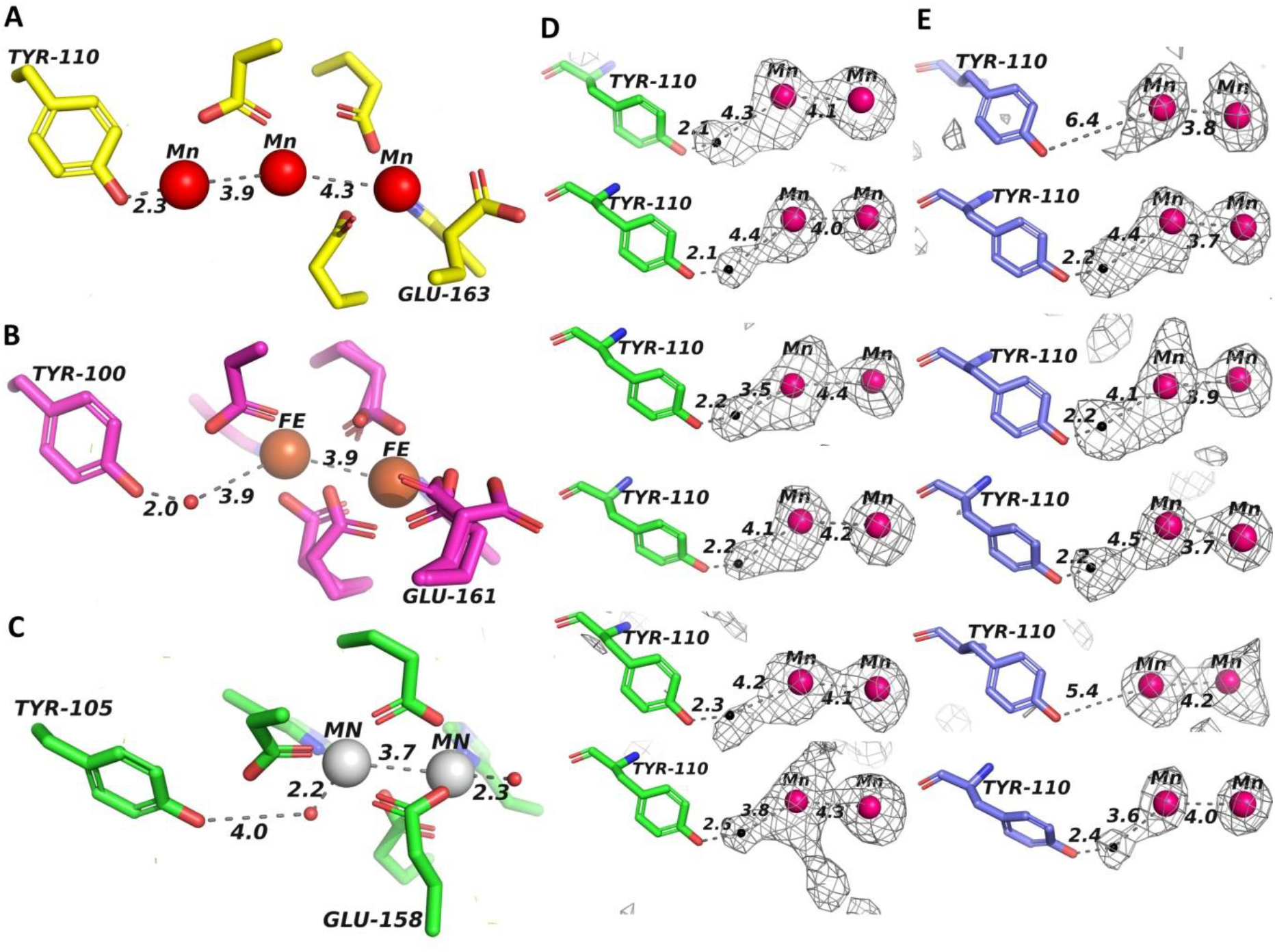
Geometry of interactions at the metal-binding site: Interaction at the metal site in NrdFI complex structures **(A)** Mtb NrdF2I**, (B)** *B.cereus* **(C).** *E.coli*. The black color sphere indicates water molecules present close to the metal cofactor site. The Fo-Fc difference density map contoured at 3σ from different NrdF2 chains of asymmetric unit in **(D)** oxidised (green) **(E)** reduced (blue) NrdF2I complex structure at metal cofactor (Mn) site.

#### 2.2.5. Binding of Mn^III^Mn^IV^ species with Tyr 110

An interesting observation in our structure is that in ten out of the twelve chains of NrdF2 (oxidised and reduced) there is a strong density between the Mn1 (Mn at site close to Tyr 110) and Tyr 110. This density is placed close to the side chain hydroxyl of Tyr 110 at a distance of 2.1-2.2Å. This distance being short for a hydrogen bond^27^ its appearance so close to the Tyr hydroxyl is perplexing. We therefore compared our structure to the reported structures of NrdF: NrdI complexes and to our surprise we found similar observation in the *Bacillus* structure also (Figure 3B and 3C). Moreover, B-factor of Mn1 in the structure is consistently higher in all the chains when compared to the second Mn-binding site (Mn2). The distance of 2.1Å is indicative of metal coordination, and therefore we believe that the density close to Tyr 110 is that of manganese ion and not that of strongly bonded water (Figure 3D and 3E). This site clearly has low occupancy as evidenced by high temperature factor. However, given the moderate resolution of the structure we did not feel appropriate to set occupancy to lower level at this site. We have therefore modelled Mn in this site.

#### 2.2.6 Tunnels to access FMN in NrdI

In the NrdF2I complex structure, a prominent tunnel is seen in NrdI starting on its surface and reaching up to FMN. Visual inspection indicates that the tunnel is constricted in the oxidised form compared to its reduced structure. In the reduced form reorientation of Arg-30 of NrdF2 and 50s loop of NrdI away from each other leads to opening of the tunnel. The tunnel radius of 2.15Å in the reduced form, we believe, is adequate for oxygen to enter and reach up to FMN (Figure 4A and B).

**Figure 4:**
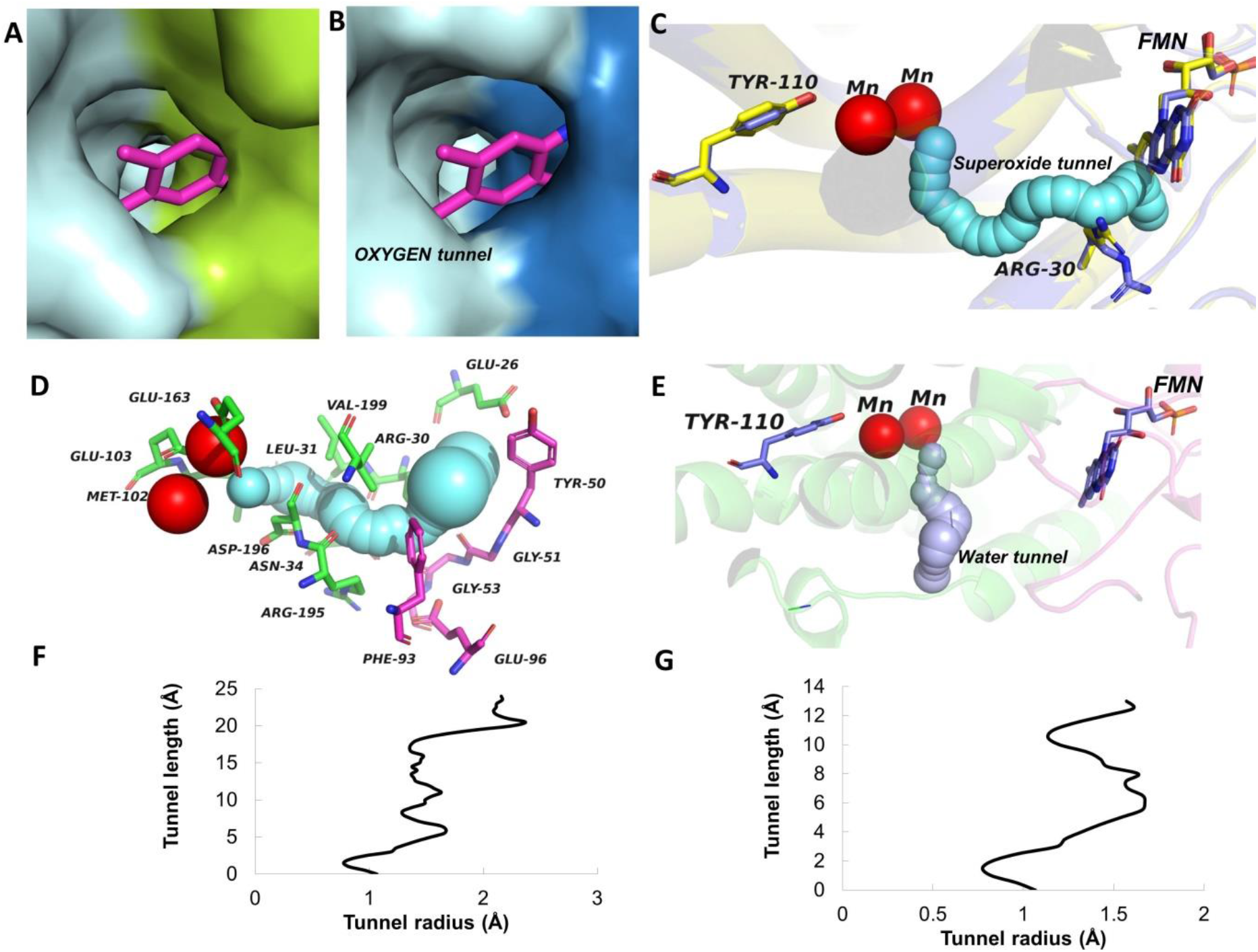
Channel formation in reduced complex of Mtb NrdF2I: Oxygen tunnel, superoxide tunnel and water tunnel connecting to FMN and metal cofactor site are important in initiation of radical formation and assembly. A prominent tunnel is seen starting from the surface of the NrdI structure and reaching FMN. We believe that this oxygen tunnel is able to carry O_2_ to the FMN site in reduced form. The oxygen tunnel is substantially constricted in the oxidized form **(A)** and expanded in the reduced **(B)** structure. **(C)** Another tunnel starting from FMN in NrdI and reaching the di-manganese site in NrdF2 is present. Formation of this tunnel is seen only in the reduced complex structure, and has been implicated in other similar structures to carry the superoxide to the metal binding site (see text for details). **(D).** Residues lining the superoxide tunnel identified by CAVER. Residues from NrdF2 are shown as green and NrdI as pink. **(E)** Water tunnel to di-manganese site in NrdF2 is present which may play important role in reduction of Mn. Profile of the superoxide tunnel **(F)** and water tunnel **(G)** indicating the radius at different parts of the tunnel.

##### 2.2.6.1 Channel for the transport of superoxide radical from FMN to the di-Manganese site

Existence of another tunnel was observed in the reduced form of the NrdF2I complex (Figure 4C). The tunnel reaches from FMN in NrdI to the dimetal site in NrdF2. The tunnel has been speculated to be the path for radical transfer to activate Mn^II^ site^26,6^. The radius of the tunnel is uniformly around 1.5Å indicating presence of sufficient space for a superoxide radical to pass through without any hindrance (Figure 4F). The reorientation of Arg 30 in the reduced complex leads to widening of tunnel gap from 1.5 to 3Å hence opening up for superoxide radical to pass through the tunnel. Of the 29 amino acids that are closer and pointing towards the 16Å superoxide tunnel are Leu-31, Arg-30, Glu-26, Val-199, Asn-34, Asp-196, Arg-195 from NrdF2 and Phe-94, Glu-97, Tyr-51, Gly 52-54 from NrdI (Figure 4D). Comparison to closest homologous *E.coli* structure indicates similar tunnel present in this complex^26^.

Interestingly a tunnel appears to be present from the surface of NrdF2 structure to the Tyr 110 residue. Although the tunnel is constricted, with minor movement of residues, the tunnel might be able to carry a water molecule to the site (Figure 4E and 4G).

## 3. Discussion

Prior studies have shown that in *Mtb* and other Mycobacteria, Class Ib NrdF2 provides the only and essential ribonucleotide reductase activity, although there are paralogs of NrdF present on the genome^11^. Different subunits of RNR, namely NrdE, NrdF2, NrdI and the thioredoxin-like protein NrdH therefore offer as attractive drug targets^28^. Structural information on these has been limited, and especially the structures of their complexes are not known. In this report we present the crystal structures of NrdI and NrdF2I in oxidized and reduced forms. We have also been able to obtain insights into the complex structures of NrdE, NrdF2 and NrdI through single particle cryo- Electron Microscopy (unpublished).

Radical formation during ribonucleotide reduction in Class Ib RNR’s occurs at the di-metal site in R2 subunit assisted by superoxide generation at the FMN site in NrdI. It has been reported that R2 or NrdF subunits of Class Ib RNR’s, when expressed heterologously in *E. coli,* are found to contain Manganese ^29^. This is unlike the equivalent NrdB subunit of the Class Ia RNR, where Iron is found to bind to NrdB. In our study on NrdF2I complexes, the identity of metal was indeed observed to be Manganese by X-ray fluorescence scan. The two Mn are coordinated, as in other reported structures, by a number of Glu, Asp and His residues^26,6^. The structures of NrdF2I complexes, in oxidized and reduced forms, thus reveal the free radical generation steps in their elegant details.

### 3.1. 50s loop of NrdI

In the atomic resolution structures of NrdI, distinct canonical features of NrdI are observed. For example, upon reduction by dithionite a peptide flip at the FMN binding site is seen, which takes place similarly in the reported structures of its homologs^19^. The peptide flip in reduced NrdI promotes strong hydrogen bonding interaction between Gly 50 carbonyl oxygen and N5 of FMN. Moreover, the interaction also stabilises the 50s loop as indicated by the B-factor plot (Supplementary Figure 2B). Unique characteristic of Mth NrdI (and that in Mtb) is the longer length of the 50s loop. This loop is disordered in the oxidized form of NrdI and assumes better order in the reduced form. Analysis of Class Ib NrdI structures, namely those from *E. coli* (3N39), *Bacillus anthracis* (2XOD), *Streptococcus sanguinis* (4N82), *Bacillus cereus* (2X2O) provide some hint to variability of 50s loop length near FMN. The organism where loop length is small (e.g *Bacillus* species), presence of Phe 45 which is part of 50s loop, acts as a cap to FMN binding^19^. Whereas in cases where the loop is longer in length, glycine residue is present at the corresponding position and the loop orients itself in a way where it covers the FMN binding site. The length of 50s loop in increasing order is of *B.cereus* (4BMO) < *E.coli* (3N39) < *A. urinae* (7MMP) and Mtb NrdI in published complex structures. In the complex structure of Mtb and *A. urinae* (Class Ie PDB ID 7MMP) the long 50s loop of NrdI is present close to NrdF2. The longer 50s loop in Mtb may provide stabilising interaction where residues 49-53 of this loop play important role in intersubunit interaction. Admittedly, rest of the residues in this loop are not seen to make much intersubunit interaction in the complex. NrdI in NrdF2I complex promotes formation of more number of hydrogen bonds and hydrophobic interaction compared to structures where length of loop is shorter.

### 3.2. Unusual stoichiometry of NrdF2: NrdI complex

One of the intriguing observations in our structures relates that to the stoichiometry of the NrdF2I complex. Stoichiometry of the complex as seen in our structure is 2:1 between NrdF2 and NrdI subunits. This is different than observed in other structures, where 1:1 stoichiometry has been reported^6,26^. Our SEC MALS study and single particle CryoEM images appear to reinforce the 2:1 stoichiometry. Further detailed studies will be needed to confirm the same.

The 2:1 stoichiometry, when viewed in the context of asymmetric NrdEF complex appears to be justified. The formation of these complexes is such that only one NrdE subunit interacts with one NrdF subunit in an asymmetric fashion. Thus, if the stoichiometry of NrdF:NrdI were 1:1, free radical would have formed in both the subunits of NrdF simultaneously, whereas only one NrdF subunit would transfer the free radical to NrdE in the NrdE:NrdF complex. We therefore believe that simultaneous formation of free radical in both the NrdF monomers facilitated by NrdI might not be required. However, binding of one NrdI subunit to one chain of NrdF2 dimer and how does it preclude binding of another NrdI molecule in order to have stoichiometry of 2:1 is not clear.

### 3.3. Solvent channel accessible to FMN and from FMN to metal cofactor site in NrdF2

One of the hitherto unknown features of the NrdF2I complex has been the mode of accessibility of oxygen to the FMN site in order to generate a superoxide radical. Our observation of a tunnel in the interaction interface with a clear access to FMN explains this elegantly. The tunnel is seen to be wide enough to carry an oxygen molecule when NrdI is in the reduced form unlike oxidized form, where tunnel diameter is constricted. It is well known that a fully reduced FMN can transfer electron to O_2_ to form a superoxide radical. We propose that as soon as oxygen is converted into superoxide radical and NrdI is oxidised, Arg-30 side chain and 50s loop move towards each other constricting the channel and setting the stage for the superoxide radical for further transition to the di-metal site in NrdF2 structure (Figure 2C and 2D). We believe that the oxidation-reduction process modulates the access of oxygen at this site such that only reduced FMN would be able to receive oxygen for the subsequent cycle of superoxide radical generation.

Another tunnel between the FMN binding site of NrdI and di-Manganese binding site of NrdF2 is observed in the reduced NrdF2I complex. Observation of this tunnel has been noted in other structures and has been speculated to carry the superoxide radical from the FMN to the di-Manganese site^26,6^. Formation of this tunnel is facilitated by subtle structural changes that occur at the interface of the NrdI and NrdF2 subunits. Prominent among these is the flip of Arg 30 side chain, which is involved in intersubunit hydrogen bond in the oxidized complex, but its flipping away from the interface in the reduced form.

### 3.4. The role of Mn^III^ Mn^IV^ in Tyr• radical formation

Elegant spectroscopic and EPR studies have highlighted the role of transient Mn^III^ Mn^IV^ species in order form Tyr• radical^15^. The transient Mn^III^ Mn^IV^ activates a water molecule to form OH^•-^ radical, which in turn can attack Tyr 110 and through a proton coupled electron transfer to form the Tyr• radical. Intriguingly in our structures, we found presence of density in ten out of twelve chains of NrdF2 close to Tyr 110 residue (Figure 3D and 3E). The density is at a very short distance from Tyr-OH, i.e. 2.1-2.2 Å, too short to form a hydrogen bond with a water molecule. Similar density at 2.1 Å is also observed in the structure of *Bacillus cereus*, but has been interpreted as a water molecule. We believe that this density might correspond to a Mn ion, and not water. Moreover, the first Mn site in its canonical binding region possesses higher B-factor than the second site. Also, in the standalone structure of Mtb NrdF2 (PDB ID 1UZR), there is a water molecule present at this site, but it is at a distance of 2.8-2.9Å from the Tyr–OH, which is a classical hydrogen bonding distance. Thus, although resolution of our structure does not permit us to make a bold statement, nonetheless we believe that most probably this density corresponds to Mn ion which shuttles between the canonical binding site and a site closer to Tyr 110. Such a displacement of Mn has been known in Photosystem II, where one of the Mn acts as the “dangler Mn”^30^. If indeed this is the case, this observation would strongly indicate the mechanism of Tyr• formation mediated directly by Mn^III^, in physical contact with each other. The redox potential and pKa of the groups involved for such a radical formation reaction might be modulated by the strongly hydrophobic environment of this site.

Thus, a channel in the reduced NrdF2I structure for carrying oxygen from surface of the molecule to the FMN site, another channel to carry superoxide from the FMN in NrdI to di-Manganese site in NrdF2, disorder in the second Mn-binding and its physical contact with Tyr 110, and a water channel to the Tyr 110 of NrdF2 would complete the structural view of radical generation in the Class Ib RNR family. An animation capturing these steps is depicted in Supplementary video1. Molecular simulations and a higher resolution view of the structures will further assist in understanding of these steps.

## 4. Materials and methods

### 4.1. Cloning of *nrdI* and *nrdF2* genes from *Mycobacterium* species

#### Restriction free cloning

The genes from the RNR operon of *Mycobacterium* species were cloned using restriction free cloning method. The subunits of RNR (NrdI and NrdF2) from the genomic DNA of Mth and Mtb were amplified using GC master mix containing Phusion DNA Polymerases. The amplified product (megaprimer) was incorporated in modified pET 32a+ vector by RF cloning method^31^. Modified pET 32a+ vector has resistance markers changed so as to facilitate co-transformation for co expression and purification. Mth NrdF2 clone in pET28a+ was a kind gift from Dr. Pratibha Tiwari and was used for co-expression and purification of NrdI. The clones were screened for presence of insert using restriction enzyme BAMH1. The sequence of clone was confirmed by sequencing.

The cloned construct had His-MBP tag with TEV protease site between the tag and the cloned gene so as to enable cleavage of the His-tagged MBP from native protein. The primers used for amplification of different subunit genes are listed in Supplementary Table 4.

The synthesis of mega primer was performed using GC rich Phusion High-Fidelity PCR Master Mix with 10µM of both primers, and 80-100 ng of genomic DNA as template in 30μl of total volume. The samples were preheated for 1 min at 98°C and then 28 cycles of PCR were performed under following condition: 20s at 98°C, 40sec at 60°C and 1 min at 72°C. After the last cycle, the reaction was incubated for an additional 10 min at 72°C. The PCR products were then purified using Qiagen gel elution kit to remove excess primers and enzymes. The megaprimers synthesized were then used as insert for 2^nd^ round of PCR along with the vector. The PCR conditions were 30sec at 98°C and then 30 cycles of PCR were performed under following condition: 10s at 98°C, 1min at 60°C and 7 min at 72°C and last cycle at 72°C for 10 min. This amplified product were then digested with DpnI to get rid of parent DNA, and transformed into *Escherichia coli DH5α* cells and positive clones were screened.

### 4.2. Purification of different subunit of RNR

#### 4.2.1. Mth NrdI

As several attempts to purify NrdI alone were unsuccessful, the protocol where NrdI co-purified with NrdF2I complex were used for purification of NrdI. *E. coli* bacterial strain *Origami 2(DE3)* was used for large scale expression and purification of Mth NrdF2I (pET28a+ /kanr-NrdF2 and Camr His-MBP NrdI) construct. Single colony was inoculated in Luria Bertani broth containing kanamycin (50 μg/ml) and chloramphenicol (15 μg/ml) and the culture was allowed to grow overnight. This grown culture was then diluted 100 fold and further allowed to grow till the OD_600_was 0.6-0.7. The culture was then induced with 0.4 mM IPTG for 14 h at 20°C. 1 litre culture pellet were harvested & resuspended in the 50 ml lysis buffer containing Tris 50mM pH 7.5, NaCl 300mM, EDTA 0.5mM and 3% glycerol. Cells were lysed by sonication at amplitude of 50, for 2 min with ON/OFF cycle of 2/15 sec. Cell debris was removed by centrifugation at 14,000 rpm for 30 min at 4°C. The protein in soluble fraction was affinity purified using amylose resin for 1 h on ice by batch method. To remove the unbound protein, resin was washed with plain lysis buffer. To obtain protein in native form, on-beads cleavage with thrombin and TEV was performed at 4°C for 18 hrs. The protein was eluted from the resin using lysis buffer. The eluted protein was passed through Ni-NTA resin to get rid of contaminating MBP tag present. The protein was concentrated and further purified using Sephacryl 16/60 120 ml column. The fractions containing purified NrdI protein was pooled, concentrated and used for further experiment.

#### 4.2.2. Co-purification of Mtb NrdF2I complex: (His-NrdF2 ampr and His-MBP-NrdI – camr)

Mtb NrdF2I construct was co-transformed in *BL21 (DE3) strain.* 800 ml culture pellet was resuspended in 40 ml lysis buffer Tris 50mM pH7.5, NaCl 300mM, EDTA 0.5mM, 3% glycerol. Sonication was done at amplitude 40, ON/OFF cycle of 2/15sec using medium size probe for 2 min. Cell debris were removed by centrifugation at 12,000 rpm at 4°C for 45 min. Ni-NTA affinity purification was done by batch method for 90 min at 4°C. Non-specifically bound proteins were removed by washing with 50 mM imidazole in lysis buffer. Protein was eluted with 500mM imidazole, 4 fractions of 5 ml each. Protein was concentrated upto 2.5 ml in 3 kDa amicon filter. Buffer exchange was performed using PD-10 column to remove imidazole. Protease digestion with TEV was performed overnight for cleavage, and His-MBP tag was removed by passing through 30 µl of Ni-NTA resin. The protein was finally concentrated upto 2 ml and injected in 16/600 Superdex 200 column. To obtain more than 90% pure protein the fraction containing NrdF2I complex was reinjected in 10/300 Superdex 200 column with FPLC buffer Tris 25mM pH 7.5, NaCl 150mM.

### 4.3. Crystallisation of Mth NrdI and Mtb NrdF2: NrdI complex

Mth NrdI (4 mg/ml) was crystallised using hanging drop vapour diffusion method. Protein concentration was estimated using nanodrop spectrophotometer at 280 nm. Crystals of NrdI were observed in PEG/Ion HT ™ in condition D5 which after optimisation were grown in mother liquor containing 0.15M potassium phosphate monobasic and 18% PEG 3350. The trials were set in ratio of 1:1 and 1:2 for protein and buffer respectively at 20°C. Diffraction quality crystals were obtained within 10 days of setting the trial. To determine the structure of reduced form of NrdI, the NrdI crystals were soaked in 800 mM sodium dithionite in mother liquor till the yellow color of NrdI crystals disappeared. The oxidised and reduced crystals of NrdI were soaked in mother liquor solution with 20% glycerol as cryoprotectant and flash frozen in liquid nitrogen.

Mtb NrdF2I complex was concentrated up to 9 mg/ml and was used for crystallisation. Protein concentration was measured using nanodrop at 280 nm. Initial crystallisation screening was performed using screens from MD, Hampton research and Jena Bioscience at 20°C in different ratio of 1:1 and 1:2 for protein and buffer respectively in 96 well MRC crystallisation plate. Of the multiple hits optimised, those obtained in Proplex (D5 and E2) and crystal screen (F9) further yielded diffraction quality crystals. The best data were obtained from crystals grown in optimised crystal screen F9 condition containing 0.1M MES pH 6, 1.4M NaCl, 0.075M sodium phosphate monobasic, 0.075M potassium phosphate monobasic. The crystals were grown to their full size in 10 days in drop ratio of 1:1 in VDX48 well plate containing 200µl mother liquor. The crystals were cryo protected using 25% glycerol in mother liquor and flash frozen in liquid nitrogen and shipped for data collection at synchrotron.

### 4.4. Data collection of Mtb NrdF2I complex and Mth NrdI

The X-ray diffraction data of Mtb NrdF2I complex (reduced and oxidised form) and NrdI (oxidised) were collected at Diamond Light source, and NrdI reduced data were acquired at ESRF_ID23 beamline. The NrdI oxidised and reduced data were acquired at 0.25 /0.25 and 0.07/0.2 exposure (sec)/ rotation (degree) respectively. Data of Mtb NrdF2I complex were collected at Diamond light source with exposure (sec)/ rotation (degree) of (0.1/0.5) and (0.2/0.2) for oxidised and reduced complex respectively. All the data of oxidised and reduced crystals were collected at wavelength of 0.9Å and structure was determined in space group of P 1 21 1 and P 42 21 2 for NrdI and NrdF2I complex respectively.

### 4.5. Structure solution refinement

Data reduction and scaling for both oxidised and reduced Mth NrdI and Mtb NrdF2I complex were performed using iMOSFLM in CCP4i^32^. The coordinates of PDB ID 1UZR and PDB ID 3N39 were selected and sequence modified in chainsaw and were then used for molecular replacement in PHASER^33^. Model building was performed in COOT^34^ and refined using REFMAC5^35^ and phenix refine^36^

For NrdI reduced, multiple data collected were merged using Blend program^37^. The cluster of merged data that had lower Rmeas and Rpim value were used for molecular replacement. Restrained refinements were performed for model building. TLS and anisotropic refinement were performed in final rounds of refinement. For NrdF2I complex, MR with PDB ID 1UZR identified 6 chains of NrdF2 in asymmetric unit. Fo-Fc density were evaluated and two molecules of NrdI was placed in the asymmetric unit by superposition of NrdF2I (PDB ID 3N39) complex structure of *E.coli* followed by fitting and rigid body refinement. Further refinement was done using REFMAC5 and Phenix refine. The TLS group for crystallographic refinement was determined using TLSMD server^38^. Structure validations of the coordinates were performed using MolProbity^39^. PyMOL (The PyMOL Molecular Graphics System, Version 1.2r3pre, Schrödinger, LLC.), Coot^34^ and Chimera^41^ were used for visual analysis and figure preparations.

### 4.6. X ray fluorescence spectroscopy

To identify the metal present in Mtb NrdF2I complex, X-ray fluorescence (XRF) scan was obtained of complex crystals. Protein crystals were washed in mother liquor with 25% glycerol and flash frozen in liquid nitrogen. XRF scan were recorded at XRD2 beamline in Elettra using X-rays at 12.3 keV and exposure time of 15 sec.

### 4.7. SEC-multiangle light scattering

120 µl of SEC purified protein at a concentration of 1.2mg/ml (Mth NrdI), 2.7mg/ml (Mtb NrdF2) and 2.7mg/ml (Mtb NrdF2: NrdI) was subjected for Size Exclusion Chromatography coupled with multi-angle light scattering (SEC-MALS). Wyatt Dawn Heleos-II light scattering detector and Wyatt Optilab T-rEX refractive index detector were connected in-line with the chromatography system for simultaneous MALS measurements. All experiments were performed in Superdex 200 10/300 GL column; (GE Healthcare) with a buffer containing 50 mM Tris pH 7.5, 150 mM NaCl at room temperature. BSA (1mg/mL) was used as standard and data was analyzed with Astra software (Wyatt Technologies).

### 4.8. Tunnel analysis

CAVER 3.0 PyMOL plugin was used for analysis and visualization of tunnels in NrdF2I complex^42^. 0.7Å minimum probe radius, 4 and 3 shell depth and shell radius respectively were parameters used for tunnel analysis. Tunnel radius versus tunnel length were plotted to determine the minimum bottleneck radius. Cavity detection radius of 7Å and cavity detection cutoff of 3 solvent radii in PyMOL were used to analyse the oxygen tunnel in both oxidised and reduced NrdF2I complex

### 4.9. PDB ID

The coordinates of Mth NrdI in reduced and oxidised form are deposited in protein data bank under PDB ID 8J4W and PDB ID 8J4V. Mtb NrdF2I complex coordinates are deposited in reduced and oxidised form in protein data bank under PDB ID 8J4Y and 8J4X.

## Acknowledgements

This work was supported by DST-NPDF (PDF/2015/000961) and DBT-Centre of Excellence Grant (BT/PR15450/COE/34/46/2016). IISER Pune Diffraction Facility and TMC ACTREC for cryooptimisation and initial screening of crystals are gratefully acknowledged. Special thanks are to Dr. Gayathri Pananghat for her useful inputs in structure determination. ESRF ID 23 and Diamond Light Source UK for X-ray diffraction data collection are gratefully acknowledged. Raghurama P Hegde XRD2 beamline - Elettra Sincrotrone Trieste for X-ray fluorescence scan is gratefully acknowledged. The genomic DNA of *M. thermoresistibile* and *M. tuberculosis* is generous gift from Dr. Sharmistha Banerjee, University of Hyderabad.

