## Supplementary Figures for "Structural insights into the initiation of free radical formation in the Class Ib ribonucleotide reductases"

### Supplementary Data

#### Supplementary figure 1: Conformational change upon reduction of NrdI

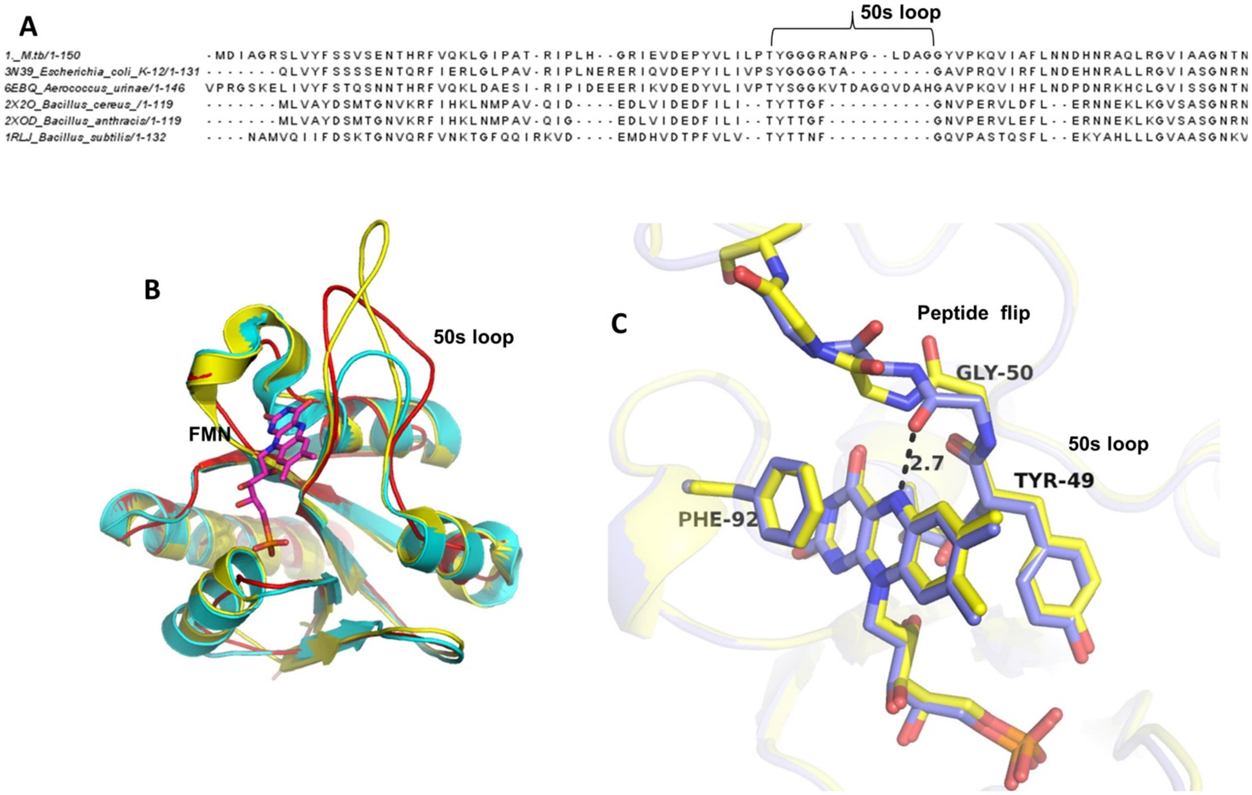


**(A)** NrdI sequence alignment from *Bacillus, Aerococcus, E.coli* and Mth shows that E. coli has a shorter 50s loop when compared with the other sequences. **(B)** Structure of *A. urinae* NrdI shown in yellow, *E. coli* in cyan, Mth in red indicates the 50s loop orients in the same direction and is the site for interaction of NrdF2. **(C) A** pronounced flip at the Gly 50 residue of 50s loop is observed in the reduced form. The peptide flip facilitates formation of hydrogen between N5 of FMN and carbonyl oxygen of the peptide.

**Supplementary Figure 2: Electron density at peptide flip and B-factor plot**

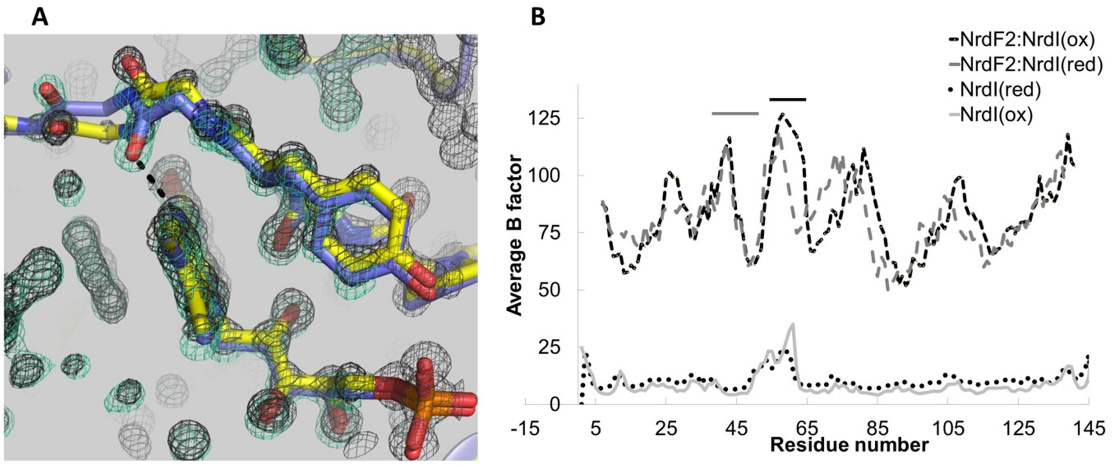


**(A)** Electron density at FMN and peptide flip in NrdI structure. **(B)** B-factor plot of NrdI in oxidised and reduced form for structure of NrdI and NrdI of NrdF2I complex. B-factor analysis of oxidized and reduced NrdI structures shows that the 50s loop gets better ordered in the latter. The grey and black line in plot indicates region of 40s and 50s loop.

**Supplementary Figure 3: Mass estimation in solution**

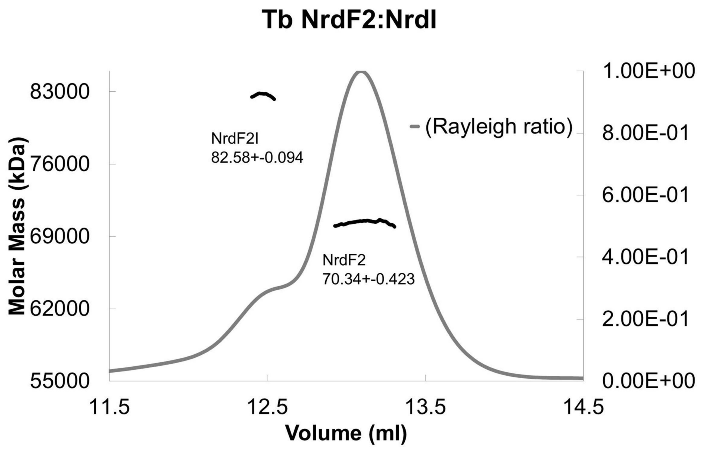


SEC MALS analysis of NrdF2I complex in Superdex 200 10/30 column.

**Supplementary Table 1:** X-ray data collection and refinement statistics.

| | **M.th NrdI(ox)*** | **M.th NrdI(red)*** | **M.tb NrdF2I(ox)*** | **M.tb NrdF2I(red)*** |
|---|---|---|---|---|
| Wavelength (Å) | 0.9 | 0.9 | 0.9 | 0.9 |
| Resolution range | 44.98 - 1.11 (1.15 - 1.11) | 45 - 1.214 (1.23 - 1.214) | 48.27 - 3.04 (3.149 - 3.04) | 86.95 - 3.02 (3.128 - 3.02) |
| Space group | P 1 21 1 | P 1 21 1 | P 42 21 2 | P 42 21 2 |
| Unit cell | 36.0 37.2 48.4 90 111.61 90 | 35.7 36.8 48.4 90 111.5 90 | 160.5 160.5 312.1 90 90 90 | 160.4 160.4 310.4 90 90 90 |
| Total reflections | 594810 (52849) | 3774695 (369168) | 1669955 (158200) | 3771275 (368185) |
| Unique reflections | 47159 (4696) | 35564 (3510) | 79042 (7798) | 80104 (7862) |
| Multiplicity | 3.9 (3.3) | 11.9 (11.7) | 11.3 (10.9) | 10.3(10) |
| Completeness (%) | 99.77 (99.68) | 99.9 (98.8) | 99.83 (99.97) | 99.90 (99.95) |
| Mean I/sigma(I) | 9.3 (3) | 9.6 (3.5) | 5.67 (1.15) | 6.3 (1.6) |
| Wilson B-factor | 7.59 | 9.2 | 54.4 | 52.58 |
| R-merge | 0.066 (0.377) | 0.143 (0.673) | 0.40 (2.15) | 0.42 (2.65) |
| R-meas | 0.086 (0.5) | 0.153 (0.736) | 0.439 (2.37) | 0.473 (2.9) |
| R-pim | 0.055 (0.325) | 0.062 (0.299) | 0.179 (0.988) | 0.204 (1.29) |
| CC1/2 | 0.998 (0.893) | 0.996 (0.94) | 0.987 (0.462) | 0.985 (0.29) |
| CC* | 0.94 (0.878) | 0.991 (0.962) | 0.916 (0.747) | 0.977 (0.749) |
| Reflections used in refinement | 47082 (4690) | 35535 (3504) | 79039 (7796) | 80102 (7858) |
| Reflections used for R-free | 2317 (252) | 1788 (158) | 3969 (426) | 4030 (436) |
| R-work | 0.1379 (0.1722) | 0.1372 (0.1753) | 0.2322 (0.3509) | 0.2310 (0.3667) |
| R-free | 0.1579 (0.2003) | 0.1710 (0.2126) | 0.2566 (0.3475) | 0.2683 (0.3875) |
| CC(work) | 0.915 (0.879) | 0.867 (0.818) | 0.819 (0.572) | 0.878 (0.534) |
| CC(free) | 0.905 (0.885) | 0.853 (0.859) | 0.837 (0.489) | 0.839 (0.523) |
| Number of non-hydrogen atoms | 1383 | 1352 | 16015 | 15947 |
| macromolecules | 1214 | 1153 | 15901 | 15852 |
| ligands | 42 | 40 | 92 | 78 |
| solvent | 127 | 159 | 22 | 17 |
| Protein residues | 145 | 144 | 1971 | 1964 |
| RMS(bonds) | 0.005 | 0.005 | 0.002 | 0.002 |
| RMS(angles) | 0.9 | 0.93 | 0.46 | 0.45 |
| Ramachandran favored (%) | 97.9 | 99.3 | 97.95 | 97.58 |

| | | | | |
|---|---|---|---|---|
| Ramachandran allowed (%) | 2.1 | 0.7 | 2.05 | 2.26 |
| Ramachandran outliers (%) | 0 | 0 | 0 | 0.15 |
| Rotamer outliers (%) | 0 | 0 | 0 | 0.06 |
| Clashscore | 0.4 | 0.85 | 2.98 | 4.17 |
| Average B-factor | 13.49 | 15 | 59.97 | 59.72 |
| macromolecules | 12.01 | 12.81 | 60.03 | 59.74 |
| ligands | 9.52 | 12 | 59.12 | 62.21 |
| solvent | 29.01 | 31.66 | 19.64 | 23.52 |
| Number of TLS groups | - | | 14 | 14 |
| *-Statistics for the highest-resolution shell are shown in parentheses. | | | | |

**Supplementary table2:** Cross Structure statistics of different structure determined of NrdI

| **Sr.no** | **Organism (PDB:ID)** | **1** | **2** | **3** | **4** | **5** | **6** | **7** |
|---|---|---|---|---|---|---|---|---|
| | NrdI | Mth NrdI | *A. urinae* 6EBQ | *B.anthracis* 2XOE | *B.subtilis* 1RLJ | *S. sanguinis* 4N82 | *D.desulfuricans* 3F90 | *E. coli* 3N3A |
| 1 | Mth NrdI | | 0.69 | 0.99 | 1.38 | 1.68 | 2.72 | 0.57 |
| 2 | *Aerococcus urinae* 6EBQ | 0.69 | | 1.05 | 1.35 | 1.57 | 2.63 | 0.76 |
| 3 | *Bacillus anthracis* 2XOE | 0.99 | 1.05 | | 0.936 | 1.49 | 2.53 | 1.061 |
| 4 | *Bacillus subtilis* 1RLJ | 1.38 | 1.35 | 0.936 | | 1.63 | 2.59 | 1.45 |
| 5 | *Streptococcus sanguinis* 4N82 | 1.68 | 1.57 | 1.49 | 1.63 | | 2.78 | 1.56 |
| 6 | *Desulfovibrio desulfuricans* 3F90 | 2.72 | 2.63 | 2.53 | 2.59 | 2.78 | | 2.71 |
| 7 | *E. coli* 3N3A | 0.57 | 0.76 | 1.061 | 1.45 | 1.56 | 2.71 | |

**Supplementary table3:** Cross Structure statistics of different structure determined of NrdFI complex

| Cross-structure statistics (RMSD Å) | | | | | |
|---|---|---|---|---|---|
| **Sr.no** | **Organism (PDB:ID)** | **1** | **2** | **3** | **4** |
| | | *B.cereus* 4BMO | *E.coli* 3N39 | *A. urinae* 7MMP | Mtb NrdF2I |
| 1 | *Bacillus cereus* 4BMO | 0 | 1.186 | 1.605 | 1.041 |
| 2 | *Escherichia coli* 3N39 | 1.186 | 0 | 0.932 | 0.631 |
| 3 | *Aerococcus urinae* 7MMP | 1.605 | 0.932 | 0 | 0.804 |
| 4 | Mtb NrdF2-NrdI | 1.041 | 0.631 | 0.804 | 0 |

**Supplementary table4:** Primer used for cloning of NrdI and NrdF2 of Mtb and Mth

| **Sr. No** | **Gene name** | **Sequence 5' - 3'** |
|---|---|---|
| 1 | M.tb nrdI F | GGATCGGAAAACCTGTATTTTCAGGGATCCATGGATATCGCGG GGCGCAGCCTG |
| 2 | M.tb nrdI R | GAACTGCGGGTGGCTCCAGCTGCCGGATCCCTACTACAGGCTCT GCAATGACGGTTGGTGG |
| 3 | M.th nrdI F | GGATCGGAAAACCTGTATTTTCAGGGATCCGTGACGGTGGGCG GCCTGGTCTAC |
| 4 | M.th nrdI R | GAACTGCGGGTGGCTCCAGCTGCCGGATCCCTATTACCGGCTCT GCAGCTGTGA |
| 5 | M.tb nrdF2 F | GGATCGGAAAACCTGTATTTTCAGGGATCCGTGACTGGAAACG CAAAGCTA |
| 6 | M.tb nrdF2 R | GAACTGCGGGTGGCTCCAGCTGCCGGATCCCTACTAGAAGTCCC AGTCATCGTCCTCGG |
| M.tb-Mycobacterium tuberculosis, M.th-Mycobacteriumthermoresistibile, F –forward primer , R –reverse primer | | |

**Supplementary video1:** The structural view and path for radical transfer and generation in the Class Ib RNR
